# Hindering memory suppression by perturbing the right dorsolateral prefrontal cortex

**DOI:** 10.1101/2024.12.23.630167

**Authors:** Davide F. Stramaccia, Frederik Bergmann, Katharina Lingelbach, Ole Numssen, Gesa Hartwigsen, Roland G. Benoit

**Affiliations:** Max Planck Institute for Cognitive and Brain Sciences, Leipzig, Germany; Institute of Cognitive Science, University of Colorado at Boulder, Boulder, CO, United States; Leipzig University, Leipzig, Germany

**Keywords:** dorsolateral prefrontal cortex, memory suppression, Think/No-Think, transcranial magnetic stimulation (TMS), retrieval stopping

## Abstract

A reminder of the past can trigger the involuntary retrieval of an unwanted memory. Yet, we can intentionally stop this process and thus prevent the memory from entering awareness. Such suppression not only transiently hinders the retrieval of the memory, it can also induce forgetting. Neuroimaging has implicated the right dorsolateral prefrontal cortex (dlPFC) in initiating this process. Specifically, this region seems to downregulate activity in brain systems that would otherwise support memory reinstatement. Here, we probed the causal contribution of the right dlPFC to suppression by combining the Think/No-Think task with repetitive transcranial magnetic stimulation (rTMS). Participants first learned pairs of cue and target words, and then repeatedly recalled some of the targets (think condition) and suppressed others (no-think condition). We applied 10 Hz rTMS bursts to the right dlPFC during the suppression of half the no-think items and to the contra-lateral primary motor area (M1) as an active control site during the other half. As hypothesized, participants experienced less success at keeping the memories out of awareness with concurrent dlPFC than M1 stimulation. Similarly, a memory test yielded evidence for suppression-induced forgetting (SIF) following M1 but not dlPFC stimulation. However, the difference in forgetting between the stimulation conditions was not significant. The study thus provides causal evidence for the role of the dlPFC in preventing retrieval. Future work will need to conclusively establish the relationship between this transient effect and suppression-induced forgetting.

## Introduction

When we are confronted with a reminder of an unwanted memory, we can prevent the memory from entering awareness (Anderson & Hulbert 2021). One mechanism supporting such mnemonic control is referred to as retrieval suppression. This mechanism is thought to avert unwanted recall by systematically inhibiting the retrieval process (Benoit & Anderson 2012; Levy & Anderson 2009). Such suppression successively weakens the avoided memory (Benoit et al. 2015; Levy & Anderson 2008; Levy & Anderson 2012), hinders the reactivation of its neural representation (Meyer & Benoit 2022), and can eventually lead to forgetting, a phenomenon referred to as suppression-induced forgetting (SIF, Anderson & Green (2001) and Stramaccia et al. (2020)).

Neuroimaging studies have provided consistent evidence that retrieval suppression is associated with increased activation in a set of brain regions that entails the right dorsolateral prefrontal cortex (dlPFC). At the same time, it is associated with reduced activation in a set of regions that are critical for memory retrieval. These most notably include the hippocampus. This pattern has been interpreted as reflecting a top-down inhibition of hippocampal retrieval processes originating from the right dlPFC (Anderson et al. 2004; Depue et al. 2007).

Over the last decade, this account has been supported by several studies that examined the functional connectivity between the dlPFC and hippocampus (Apšvalka et al. 2022; Benoit & Anderson 2012; Benoit et al. 2015; Benoit et al. 2016; Depue et al. 2007; Gagnepain et al. 2014; Leone et al. 2022; Mary et al. 2020; Paz-Alonso et al. 2013). These studies varied in procedural details, including the type of memories examined and the methods used for their initial acquisition. Yet, they consistently provided evidence that the dlPFC modulated activity in the hippocampus during suppression.

However, neuroimaging studies can only provide correlational evidence for the involvement of the right dlPFC. Here, we tested its causal contribution by complementing a Think/No-Think task (Anderson & Green 2001) with transcranial magnetic stimulation (TMS). We first had participants associate pairs of cue and target words. In the subsequent suppression phase, we then presented some of the cues repeatedly in green font and requested participants to covertly recall the associated target (think items). We presented other cues repeatedly in red front (no-think items) and instructed participants to prevent the associated targets from coming to mind. In case a memory intruded into their mind, they were instructed to push it back out of awareness. We thus used instructions that have previously been associated with the typical activity pattern of retrieval suppression (Benoit & Anderson 2012; Bergström et al. 2009).

To assess the causal contribution of the right dlPFC, we perturbed activity in this region using 10 Hz repetitive TMS (rTMS) bursts for half of the suppress items (no-think^dlPFC^ items). For the other half of these items, we perturbed the contralateral M1 as an active control site (no-think^control^ items). We could thus test whether stimulating the right dlPFC hinders participants’ ability to prevent unwanted retrieval.

Afterward, we assessed participants’ memories of the targets from the *think* and the two *no-think* conditions (dlPFC, M1 control) as well as from a baseline condition. This latter condition comprised memories that had not been cued during the preceding suppression stage. This allowed us to infer whether suppression exerts an impact that goes beyond the passive forgetting over time occurring for the baseline items. Finally, after the experiment, we measured the experienced disruption in mnemonic control by asking participants how often each memory had inadvertently intruded into awareness during no-think trials.

Based on the hypothesis that the right dlPFC - but not M1 - is causally involved in suppression, we had two predictions. First, perturbing activity in the right dlPFC should make it more difficult to prevent the retrieval of unwanted memories than perturbing activity in the active control site. Second, impeding suppression should also prevent the typical inhibitory after-effects: we expected suppression-induced forgetting only in the control condition but not for memories whose suppression had been hindered via dlPFC stimulation.

## Methods

### Participants

Thirty-one participants (23 female, 8 male; mean age = 25.29, *SD* = 4.58) took part in the experiment. They were German native speakers, right-handed, and had graduated from high school. Furthermore, they reported no color blindness, had normal or corrected-to-normal vision, and were naive to the Think/No-Think task. One participant was excluded due to technical issues, leaving a sample of 30 (23 female, 7 male; mean age = 25.37, *SD* = 4.64). We recruited the participants from the database of the Max Planck Institute for Human Cognitive and Brain Sciences, Leipzig.

Our sample size (n = 30) is common for studies examining memory suppression with the Think/No-Think procedure (24–30 participants). It provides 80% power to detect a medium effect size (i.e., Cohen’s *d* = 0.47, one-tailed; Cohen’s *f* = 0.43 in a 2×3 repeated measures ANOVA) at α =0.05 (according to G*Power 3.1). This is below the metaanalytic effect-size estimate (*d* = 0.66) of SIF in healthy individuals (Stramaccia et al. 2020).

A study physician certified their eligibility for MRI and TMS, which was also assessed by the experimenters prior to each experimental session with a safety questionnaire. The study protocol was approved by the local research ethics committee (protocol number 506/16-ek) and was in accordance with the principles of the Declaration of Helsinki. At the beginning of both experimental sessions, we obtained written informed consent. All participants were reimbursed for their time.

### Think/No-Think Procedure

Our Think/No-Think procedure comprised three phases (Figure 1): First, a study phase, in which participants learned cue-target word pairs; second, a Think/No-Think phase, in which a subset of cues were shown again: some cues in red, indicating that the target is to be suppressed (*no-think*), and some in green indicating the target is to be retrieved (*think*). And third, a test phase, in which they attempted to recall all targets.

**Fig. 1.**
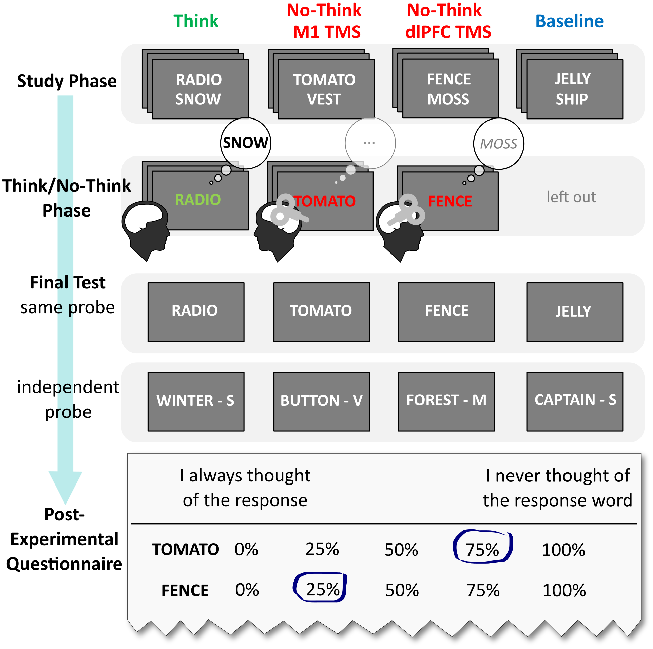
Experimental Procedure. In a study phase, participants learned associations between pairs of cue and target words (stimuli are translated from the original German for illustration), without knowing that the pairs would appear in different conditions later. In the subsequent Think/No-Think phase, they recalled target words prompted with green cues, whereas they attempted to suppress the retrieval of targets prompted with red cues. During these trials, we either stimulated the left M1 (as an active control site) or right dlPFC. Targets from the Baseline condition were not cued during this phase. They thus serve as a baseline measure of memory performance on the two final memory tests. After the final recall tests, participants filled out a post-experimental questionnaire. Here, they were given a list of all No-Think cues and judged to what degree they were able to keep the associated targets out of mind during the TNT task.

### Material

Stimuli consisted of 50 German cue-target word pairs and another word that was semantically related to the respective target (e.g., RADIO – SNOW – WINTER). The latter word served as an independent memory probe on the final memory test (see below). Most of the stimuli were translations from a Dutch set (van Schie & Anderson 2017). They were divided in five sets of 10 pairs. An additional set of 8 stimuli was used to practice the tasks and to minimize primacy and recency effects during the study phase. The five critical sets were comparable in terms of word frequency as well as orthographic and phonological properties as indexed by the dlexDB (Heister et al. 2011) and the Leipzig Corpora Collection (Goldhahn et al. 2012).

We adopted a Latin-square counterbalancing strategy so that, across the sample, each item appeared equally often in all conditions (*baseline, think, no-think*) and stimulation sites (dlPFC, M1). For each counterbalancing condition, one set of stimuli was assigned to *baseline* items, whereas *think* and *no-think* items were assigned two sets each. Participants thus encountered as many *think* as *no-think* items in the Think/NoThink phase. This approach led to ten counterbalancing conditions, which we repeated three times across the 30 participants.

During each phase of the procedure, we ensured that (a) a maximum of three items of the same condition were presented consecutively, (b) the same item could reappear only once all other items had been presented, and (c) the average serial position of the items was similar for all conditions.

### Study Phase

In the initial study phase, participants encoded 50 critical cue-target pairs. Participants also learned a further 8 filler pairs, which were used during training the task and to mitigate primacy and recency effects during learning. First, we presented half the critical and filler pairs for 4000 ms each, followed by a fixation cross for 400 ms. Afterward, we assessed participants’ memory for the pairs in a series of test-feedback cycles. Here, participants encountered each cue word for 3500 ms and attempted to covertly recall the associated target. Irrespective of their response, the correct target was then displayed in blue font for 1000 ms. Each trial started with a fixation cross for 1000 ms. Participants carried out these test-feedback cycles until they had correctly recalled at least 50% of the targets in a single run. This procedure was then repeated for the second half of the word pairs. Afterward, participants once more attempted to recall each target in response to its cue, which was presented for 3500 ms, each trial preceeded by a fixation cross for 1000 ms. No feedback was provided at this stage. On average, participants were able to learn about 75.7% of the stimuli by the end of the initial study phase (*SD* = 11.9%).

### Think/No-Think Phase

In the following Think/No-Think phase, forty cue words were presented one at a time for 2500 ms, either in green (*think* items, n = 20) or red color (*no-think* items, n = 20). Each trial started with a fixation cross for 400 ms and was followed by a blank ISI of 400 ms. For the green *think* items, participants were instructed to recall the associated target silently and to rehearse it in their mind for as long as the cue remained on screen.

For red *no-think* items, participants were instructed to avoid thinking about the associated target at all costs while focusing on the presented cue. If a target intruded into their awareness, they were supposed to push it out. Furthermore, they were asked to keep doing so during the subsequent fixation period. We thus used instructions previously shown to induce a mechanism of retrieval suppression that presumably involves an engagement of the right dlPFC (Benoit & Anderson 2012).

After receiving the instructions, participants practiced the tasks on the filler items in two blocks. They then entered the Think/No-Think phase proper. During this phase, each critical cue was repeated ten times for a total of 400 trials across all conditions, divided into five blocks. Short breaks were provided between blocks, which also allowed us to adjust the positioning of the TMS coils if necessary. Furthermore, brief questionnaires were administered after both practice rounds and after the third experimental block to ensure that they were following the instructions to engage in retrieval suppression.

### Final Test Phase

In the final phase, we tested memory for all 50 targets using two different tests. First, on a same probe (SP) test, we presented the original cue, and participants attempted to recall the corresponding target aloud. Second, on an independent probe (IP) test, we probed the memories with novel words that were not previously shown during any phase. These words were strongly associated with the respective target only. They were presented together with the target’s first letter (for example, WINTER S… to test memory for SNOW). The task was to say aloud the target word with the given starting letter that matches this novel cue. The IP procedure has been designed to test whether suppression weakens the very memory trace of the target rather than merely the association between cue and target (Anderson 2003). On each test, a trial started with a fixation cross of 1000 ms followed by the cue shown for 3500 ms. Participants practiced both tests on the filler pairs, and they always received the SP test first.

### Apparatus

In the study and final test phase, words were presented in white (cues) and blue (targets) on a gray background, on a 23-inch color flat screen display (EIZO EV2450 FlexScan) with a maximal resolution of 1920 × 1080 pixels and a refresh rate of 60 Hz. During the Think/No-Think phase, stimuli were either presented in green or red on a gray background on a 17-inch color flat screen display (EIZO FlexScanS1901) with a maximal resolution of 1280 × 1024 pixels and refresh rate of 60 Hz.

Participants were seated ~50 cm (study and final phase) and ~125 cm (Think/No-Think phase) from the screen in a quiet, dimly lit room. Words were presented at the center of the screen, with a letter width and height of approximately 1.3 cm for the shorter and 3 cm for the longer viewing distance (equivalent to a visual angle of 1.49° and 1.38°, respectively). PsychoPy version 1.85.3 (Peirce 2008) was used to run the experiment and to control the TMS pulses via parallel port.

### Transcranial magnetic stimulation

#### Apparatus and Target Localization

We delivered 10 Hz rTMS bursts with two C-B60 figure-of-eight coils attached to two MagVenture MagPro X 100 stimulators. We used two coils to alternate stimulation over the right dlPFC and the M1 control site, thus allowing us to test our main hypothesis in a within-participant design. To ensure precise coil positioning, we used a neuronavigation system (TMS Navigator, Localite GmbH, 2015) with individual T1-weighted MRI scans. These were acquired prior to the experiment with a three Tesla Siemens Magnetom Prisma Scanner (Siemens Healthcare GmbH, 2018). The T1 scans were converted to DICOM format und then segmented and normalized using SPM8. After visual inspection the target coordinates (see below), they were then transformed from MNI to native space using the SPM8 function spm_get_orig_coord.

The target position for each coil was based on the MNI coordinates obtained from Benoit & Anderson (2012) for the right dlPFC, and from a meta-analysis by Mayka et al. (2006) for the left hand area in M1. Specifically, for the right dlPFC, we used the group-level peak coordinates from the contrast associated with retrieval suppression (i.e., Suppress > Recall, x = 36, y = 38, z = 31). For M1 we chose the hand-responsive area, defined by the peak activation-likelihood es-timate across motor tasks involving the right hand (x = −37, y = −21, z = 58). We converted the MNI coordinates to each participant’s MRI native space and overlaid them on their individual structural scans. Once in place, we manually maintained the coils’ position during the experiment to enable onthe-fly adjustments to participants’ movement. Because the neuronavigation system could only display a single coil at a time, we tracked and maintained the position of the dlPFC coil online to maximize the precision of the critical stimulation and only adjusted the position of the control M1 coil during the breaks between blocks of the task.

We oriented both coils at 45° relative to the mid-sagittal axis (with handles pointing backward) to induce a posterior-anterior current flow approximately perpendicular to the central sulcus. Due to technical limitations of the neuronavigation system, we were only able to continuously monitor the movement of the right dlPFC coil throughout the experimental blocks. The position of the M1 coil was checked during each break and adjusted accordingly.

#### Stimulation Parameters

We employed a 10 Hz repetitive TMS stimulation protocol which is thought to lead to a suppression of activity during the first 100 to 200 ms after a TMS pulse Silvanto & Pascual-Leone (2008). Meta-analytic evidence has corroborated that rTMS at a frequency range from 10 Hz and 20 Hz can disrupt attention, executive function, language, memory, motor, and perception Beynel et al. (2019). As such, this protocol has been successfully used for perturbing higher cognitive functions in event-related designs similar to the one employed in the current study (e.g. Kennerley et al. (2004), Kuhnke et al. (2017), Nixon et al. (2004), and Sandrini et al. (2008)). Importantly, this protocol allowed us to stimulate the respective target areas during all 400 no-think trials while complying with standard safety guidelines (Rossi et al. 2009; Rossini et al. 2015).

Specifically, on each trial, we delivered bursts of five 10 Hz rTMS pulses (100 ms inter-pulse interval) that lasted for 400 ms. For each of the two stimulation sites, this resulted in a total of 500 TMS pulses. We used an intertrial interval of 3 s to minimize (i) carry-over effects of stimulation from one trial to another, (ii) and summation of excitability due to trans-synaptic communication, while (iii) staying within safety limits (Rossi et al. 2021). Our interstimulus interval of three seconds has previously been shown to prevent any spillover effects from one trial to another (Hamidi et al. (2011) see also Julkunen et al. (2012)). Indeed, the effective gap between stimulations was even longer in this study, because the suppress trials alternated with recall trials during which we did not apply any stimulation.

Within each *no-think* trial, we started the stimulation 300 ms after the red cue had appeared on the screen, at a time when its sensory and linguistic properties could already have been processed. Critically, in doing so, we targeted a time window that likely marks the earliest deployment of inhibitory processes in the service of memory suppression as indicated by the emergence of a frontal negative eventrelated potential (Hellerstedt et al. 2016). We thus assumed that our stimulation would interfere with the suppression process early on.

We adapted the stimulation intensity based on each participant’s individual resting motor threshold (rMT), defined as the minimal intensity required to elicit a motor-evoked potential (MEP) of at least 50 µV in five out of ten consecutive TMS pulses to the primary motor cortex (Rothwell et al. 1999). In an intake session, prior to the main experimental session, we measured the rMT with single TMS pulses (0.2 Hz) for the right first dorsal interosseous muscle.

To measure the individual resting motor threshold, electromyography (EMG) electrodes were connected to a patient amplifier system (D-360, DigitimerLtd., UK, Welwyn Garden City; bandpass filtered from 10 Hz to 2 kHz), which in turn was connected to a data acquisition interface (Power1401 MK-II, CED Ltd., UK, Cambridge, 4 kHz sampling rate). Electromyography was recorded using the Signal software (CED, version 4.11).

Adopting a conservative approach, we began stimulating at a 35% intensity and gradually increased intensity in steps of 1% to 3% while systematically searching for the coil position, tilt, and orientation that most reliably generated MEPs. We adjusted each participant’s individual stimulator intensity to 90% rMT for the subsequent experimental session (Rossini et al. 2015). The average stimulation intensity resulting from this procedure was 41.3% of maximum stimulator output (*SD* = 5.06, range: 30% to 54%).

#### Questionnaires

In the intake session, participants completed the Thought Control Ability Questionnaire (TCAQ; Luciano et al. (2005)), *M* = 84.56, *SD* = 19.62, range: 25-113, and the State-Trait Anxiety Inventory (STAI; Laux et al. (1981) and Spielberger et al. (1970)), using both the state subscale (STAI-S): *M* = 32.17, *SD* = 6.59, range: 21-45; and the trait sub-scale (STAI-T): *M* = 35.8 *SD* = 6.94, range: 25-51.

Following the Think/No-Think procedure, we also assessed the impact of stimulating the right dlPFC on people’s ability to control unwanted retrieval. For this, participants were given a list with all cues from the No-Think condition. For each cue, they indicated the proportion of trials on which they had successfully kept the target out of mind, using a scale ranging from 0% to 100% in steps of 25%. A lower score corresponds to a greater difficulty in suppressing the respective memory. We calculated and compared the average response for all items in the dlPFC and in the M1 conditions. The questionnaire did not indicate which item belonged to which experimental condition. Other studies have more directly assessed the difficulty of suppressing a memory by having people rate, on a trial-by-trial basis, whether a memory had involuntarily intruded into their awareness (Benoit et al. 2015; Gagnepain et al. 2017; Legrand et al. 2022; Levy & Anderson 2012). We opted for an offline measure, however, to avoid potential confounds in the interpretation of any TMS effects. For example, rating differences in the M1 versus dlPFC conditions may have reflected stimulation-induced changes in meta-cognitive operations rather than in memory suppression.

In addition, the post-experimental questionnaire served to gauge whether participants thoroughly followed the task instructions and to which extent they were distracted by the TMS at either one of the two stimulation sites. Specifically, using open questions, it asked participants whether they had experienced any distracting muscle twitches and, if so, which parts of their body had been affected. If participants reported twitches around the right eye or temple, we coded this to have occurred as a consequence of right dlPFC stimulation. If they reported twitches of the right arm or hand, we coded this as having occurred due to left M1 stimulation. In cases where a participant reported both but did not clearly state which one was more distracting, we counted this as *both*. When in doubt, we conservatively coded greater distraction to have occurred as a consequence of right dlPFC stimulation. If the participants had provided no response to these questions, we coded that *neither* of these stimulations had induced twitches.

#### E-field Modeling

Each TMS pulse generates a short and strong electromagnetic field (E-field) that penetrates the skull and cortical tissue. To precisely quantify the individual stimulation exposure in the relevant cortical areas, we computed the respective induced electric fields using SimNIBS v3.1 (Saturnino et al. 2019). First, we generated individual head models from structural MR images using the headreco pipeline (Puonti et al. 2016), employing SPM12 and CAT12 (Dahnke et al. 2013; Gaser et al. 2024). T1 images and, where available, T2 images were used for segmenting scalp, skull, grey matter (GM), white matter (WM), cerebrospinal fluid (CSF), and eyes. The final head models were composed of~1.7 × 106 nodes and~ 9.5 × 106 tetrahedra. We then defined the position and orientation of the coil for each participant and condition, using the stored *InstrumentMarker* of the Neuronavigation System (TMS Navigator, Localite) (Numssen et al. 2021; Weise et al. 2022)).

Second, we calculated the induced electric fields for 1 A/µs stimulation intensity and subsequently scaled the fields to the individual, realized stimulator intensity. Default isotropic conductivity values were used. We visually assessed the quality of the head reconstructions and electric field simulations.

Third, we mapped the electric field of each participant per session to fsaverage space for group analyses (Fischl et al. 2008). For one participant, fields could not be computed because the meshing procedure failed.

Finally, we extracted the overall field strength, i.e., the field magnitude |E|, at the cortical target. Specifically, we included all elements within a sphere (radius = 5 mm) centered on the cortical target, masked to grey matter only. We transformed the group coordinates into the individual space and, if outside of grey matter, shifted these individual coordinates to the closest grey matter.

#### Statistical Analysis

We first examined the impact of stimulating the right dlPFC on the ease with which participants were able to prevent unwanted retrieval. We therefore compared their aggregated ratings for the *no-think^dlPFC^* and the *no-think^control^* conditions. Given the ordinal nature of the responses, we used a Wilcoxon signed-rank test. Because one participant did not provide these ratings, this analysis is based on *n* = 29.

We then examined whether rTMS also attenuated suppression-induced forgetting by comparing recall accuracy between the *baseline, no-think^dlPFC^*, and *no-think^control^* conditions on the SP and IP tests. Specifically, we conducted a repeated measures ANOVA with *suppression status* (*nothink*^*dlPFC*^, *no-think*^*control*^, *baseline*) and *test format* (SP, IP) as within-subject factors. We followed up on these results with reduced ANOVAs, each of which compared two levels of *suppression status*. Follow-up t-tests were conducted if indicated by a significant interaction term.

For exploratory purposes, we also report a repeated measures ANOVA with *rehearsal status* (*think, baseline*) and *test format* (SP, IP) as within-subject factors. This analysis allowed us to assess the effects of repeatedly rehearsing a memory in the *think* condition. We carried out all analyses using JASP (JASP Team 2024).

## Results

### rTMS affects the regions of interest

The individual field modeling corroborated the effective stimulation of the two targeted sites (i.e., right dlPFC and left M1). At the same time, the stimulation did not affect the respective other cortical site (see Figure 2A for whole brain field estimates for an example subject). The average peak stimulation strength was similar for both targets (M1: 46.8 V/m ± 12.8 V/m; dlPFC: 73.7 V/m± 16 V/m (*mean* ± *SD*)). Paired-samples t-test revealed that this difference was statistically significant, *t*(28)= −8.14, *p* < .001. However, such a difference remains well within the range typically observed in TMS studies (Numssen et al. 2024). This disparity likely results from differences in cortical depth and gyrification patterns (Numssen et al.2023).

**Fig. 2.**
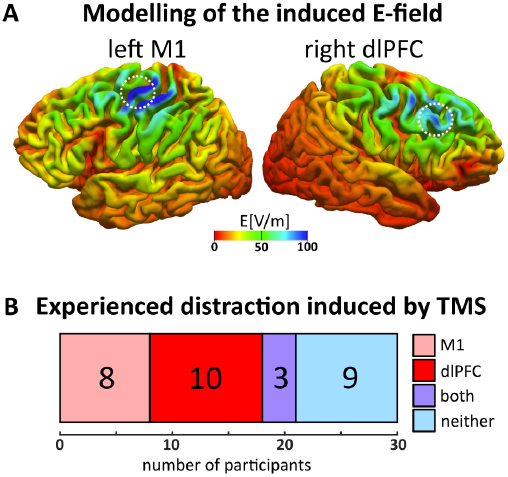
**A)** Anatomically specific stimulation of the cortical targets. Estimates for the TMS-induced E-field for left M1 and right dlPFC for an example participant. The induced field peaked at the cortical targets (dotted white circle). **B)** M1 and dlPFC stimulation appear to be similarly distracting. Participants indicated that M1 and dlPFC stimulation led to distracting muscle twitches in a similar number of participants during the experiment (shown as proportion of bar graph), indicating that this factor does not constitute a potential confound.

### M1 control and dfPFC stimulation are similarly distracting

To address a potential confound, we assessed whether either right dlPFC or left M1 stimulation may have been perceived as more distracting. Specifically, we asked participants whether they had experienced any muscle twitches induced by TMS and, if so, at which location. This analysis revealed that 9 out of 30 participants did not report any muscle twitches (Figure 2B). Three reported twitches during stimulation at both sites (*n* = 3). The remaining 17 were almost evenly split, with only two more participants reporting twitches only following stimulation of the dlPFC (*n* = 10) than of M1 (*n* = 8). This pattern suggests that the experimental effects are unlikely to be accounted for by this factor.

### Stimulating the right dlPFC makes it harder to stop retrieval of unwanted memories

Analyses of participants’ reported experience corroborated that, on average, they were less successful at not thinking about targets in the *nothink*^*dlPFC*^ condition (73.0% of the trials, score of *M* = 2.92, *SE* = 0.13) than in the *no-think^control^* condition (76.3% of the trials, score of *M* = 3.05, *SE* = 0.11), *W* = 93, *z* = 2.10, *p*= .019, one-tailed; *rank-biserial correlation* = 0.45. Thus, a perturbation of the likely source of the inhibitory process renders suppression more difficult.

### Suppression-Induced Forgetting only following stimulation of the M1 control site

We next sought to determine whether perturbing dlPFC activity on No-Think trials also affects suppression-induced forgetting (see Figure 3B). Specifically, we expected SIF for the *no-think^control^* but not the *nothink*^*dlPFC*^ condition. Our initial ANOVA with the factors *suppression status* (i.e., *no-think^dlPFC^, no-think^control^, baseline*) and *test format* (SP, IP) yielded both significant main effects *(suppression status, F*(2, 58) = 3.39, *p* = .041, η^*2*^_*p*_ =.105; *test format F*(1,29) = 6.85, *p* = .014, η^*2*^_*p*_ = .191), with higher retrieval scores on the SP than the IP test. There was no evidence for an interaction between the two factors, *F*(2, 58) = 0.14, *p* = .871, η^*2*^_*p*_ = .005.

**Fig. 3.**
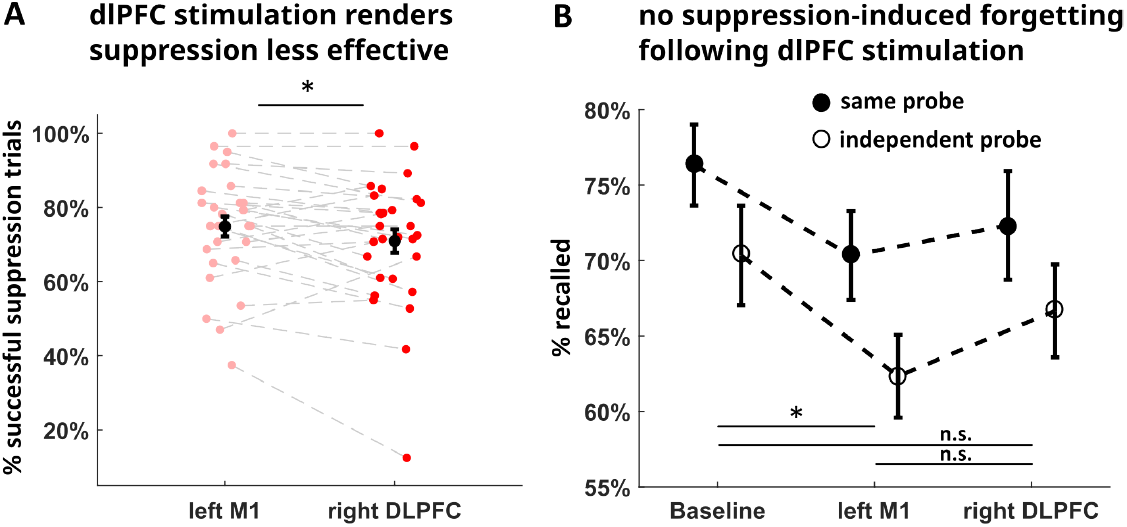
**A)** Participants experienced suppression as more difficult during stimulation of the right dlPFC than left M1 as indicated by a lower percentage of trials on which they were successful at keeping the target memory out of awareness. Black dot indicates mean and error bars standard error of the mean. **B)** The final tests yielded significant suppression-induced forgetting (lower recall for no-think compared to baseline) only for memories that had been suppressed with concurrent stimulation of left M1 but not right dlPFC. However, the interaction between suppression status and test format was not significant. Bars indicate standard error of the mean.

The follow-up analysis based on *no-think^control^* and *baseline* items indeed provided evidence for worse recall in the former condition (*F*(1, 29) = 6.63, *p* = .015, η^*2*^_*p*_ = .186). Again, *test format* was also significant (*F*(1, 29) = 7.57, *p* =.010, η^*2*^_*p*_ = .207), reflecting higher recall rates in the SP than IP test. The interaction between the two was not significant (*F*(1,29) = 0.24, *p* = .625, η^*2*^_*p*_ = .008).

By comparison, the analysis comparing *nothink*^*dlPFC*^ and *baseline* showed a trend for the effect of test format (*F*(1, 29) = 4.05, *p* = .053, η^*2*^_*p*_ = .123) with no evidence of suppression-induced forgetting (*suppression status: F*(1, 29) = 2.10, *p* = .158, η^*2*^_*p*_ = .068) or the interaction (*F*(1, 29) < .001, *p* = .944, η^*2*^_*p*_ < .001).

A direct comparison of the two no-think conditions was not significant (s*uppression status: F*(1, 29) = 1.36, *p* = .253, η^*2*^_*p*_ = .045). This analysis yielded a significant effect of *test format* (*F*(1, 29) = 4.82, *p* = .036, η^*2*^_*p*_ = .143) but no inter-action *F*(1, 29) = 0.18, *p* = .675, η^*2*^_*p*_ = .006). Though we did observe evidence for SIF in the control condition but not following dlPFC stimulation, we did not obtain significant evidence for a difference between the two conditions.

Given the limited number of cue-target pairs for each condition, the foregoing analysis was based on the average performance for all items. We thus did not exclude items that participants had not recalled at the end of the initial learning phase. Excluding those items leads to strong ceiling effects, particularly on the SP test. Nonetheless, repeating the analyses restricted only to items learned at the beginning yielded a similar pattern of results: A trend for the main effect of SIF for the control condition (*F*(1, 29) = 2.92, *p* = .098, η^*2*^_*p*_ =.091), an absent effect following dlPFC stimulation (*F*(1, 29)= 1.51, *p* = .229, η^*2*^_*p*_ = .049), but no significant effect between the two suppression conditions (*F*(1, 29) = 0.137, *p* =.714, η^*2*^_*p*_ = .005).

### Exploration of individual differences

For exploratory purposes, we correlated the questionnaires (TCAQ, STAI-S, STAI-T) with SIF on both the IP and SP tests, as well as the intrusion control rating using Spearman rank correlations. None of the correlations exceeded significance (p > .05), with rho < |.25| (see also Meyer & Benoit (2023)).

### Comparison of the Think versus Baseline conditions

Finally, we explored whether repeatedly rehearsing Think items led to better memory on the final test compared to Baseline, that is, an effect of retrieval practice (Chan et al. 2006; Karpicke & Blunt 2011; Karpicke & Roediger 2008; Roediger & Butler 2011). The corresponding ANOVA yielded a significant effect of *test format* (*F*(1, 29) = 31.86, *p* < .001, η^*2*^_*p*_ = .523) and a significant interaction (*F*(1, 29) = 6.12, *p*= .020, η^*2*^_*p*_ = .174) with no evidence for the main effect of*rehearsal status* (*F*(1, 29) = 2.64, *p* = .115, η^*2*^_*p*_ = .083).

Follow-up analyses revealed comparable recall in the *think* and the *baseline* condition on the SP test (*t*(29) = −0.25, *p* = .807, Cohen’s *d* = −.05), with only a numerical advantage for the former condition. The absence of a significant effect of retrieval practice likely reflects a ceiling effect, given the high performance observed across all conditions.

On the IP test, the *think* condition yielded worse performance than the *baseline* condition (*t*(29) = 2.54, *p* = .017, Cohen’s *d* = .46). This reversed practice effect on the IP test has already been reported in the literature (e.g., Benoit & Anderson (2012)), and it has been argued to reflect encoding-specificity effects (e.g., Paz-Alonso et al. (2009)). Given the intensive test-feedback procedure used in the study phase, we may have very well increased the likelihood of these effects.

## Discussion

This study set out to determine the causal contribution of the right dlPFC to the suppression of unwanted memories. Previous neuroimaging evidence has assigned a key role to this region in preventing the intrusion of unwanted memories. A stronger engagement of the right dlPFC leads to a steeper decline in the frequency with which unwanted memories intrude into awareness (Benoit et al. 2015), and it has also sometimes been associated with stronger suppression-induced forgetting (Anderson et al. 2004; Hanslmayr et al. 2010).

Here, we complemented the correlational evidence from neuroimaging by testing for the necessary involvement of the right dlPFC in memory control. We disrupted the suppression process via a 10 Hz rTMS stimulation over the right dlPFC and included, as an active control site, stimulation over contralateral M1. Critically, our design allowed us to assess the contribution of the dlPFC to two facets of suppression: (i) people’s transient ability to prevent unwanted retrieval and (ii) the sustained *aftereffect* of this process, i.e., suppression-induced forgetting.

The right dlPFC has been suggested to transiently prevent unwanted memory intrusions by eliciting a targeted inhibition of the hippocampus (Anderson et al. 2004) and of other regions involved in encoding the memory’s neural trace (Benoit et al. 2016; Depue et al. 2007; Gagnepain et al. 2014; Meyer & Benoit 2022). The right dlPFC would thus contribute to the reduction of a memory’s intrusiveness (Benoit et al. 2015; Levy & Anderson 2012), hinder its subsequent neural reactivation and (Meyer & Benoit 2022), reduce its vividness (Meyer & Benoit 2022) and eventually cause forgetting (Anderson & Hulbert 2021; Stramaccia et al. 2020).

This model of top-down modulation has been supported by several neuroimaging studies that reported stronger functional connectivity between the dlPFC and hippocampus during suppression. Using dynamic causal modeling, some of these provided specific evidence for a directed modulation of hippocampal activity originating from the dlPFC (Apšvalka et al. 2022; Benoit & Anderson 2012; Benoit et al. 2015; Gagnepain et al. 2014; Hu et al. 2017; Mary et al. 2020). At the structural level, this top-down modulation could be accomplished via excitatory pathways from the prefrontal cortex onto inhibitory neurons in the entorhinal cortex or via the nucleus reuniens (Anderson et al. 2016).

In the present study, we examined whether perturbing activity in the right dlPFC makes it more difficult to suppress unwanted memories. On average, participants reported that they were less successful at preventing memories from entering awareness during the targeted stimulation of the right dlPFC than of the active control site. We thus obtained evidence that the right dlPFC causally contributes to transiently preventing the retrieval of unwanted memories.

One potential confound to consider is whether dlPFC stimulation may have caused greater discomfort and thereby may have distracted participants from deploying. We addressed this question by asking whether they experienced any TMS stimulation as distracting, especially due to possible muscle twitches, and, if so, which stimulation site they felt was more distracting. Their responses did not indicate a stronger impact of stimulating either site. However, future studies may more systematically assess any physiological sensations induced by the two stimulation conditions.

This distraction measure, however, did not allow us to isolate the precise TMS side effects at each stimulation site such as the overall pain or intensity of muscle twitches. Given that prefrontal stimulation may be experienced as more aversive, future studies may want to assess these possible effects in more detail.

Our evidence for an effect of right dlPFC stimulation is less conclusive with respect to the sustained aftereffects. We indeed only observed suppression-induced forgetting for memories that people tried to suppress with concurrent M1 but not dlPFC stimulation. However, the critical comparison of these stimulation conditions was not significant. We thus did not obtain direct evidence that dlPFC stimulation leads to reduced SIF compared with M1 stimulation.

There are several potential reasons for this absence of evidence. First, it is conceivable that the dlPFC only contributes to the momentary prevention of unwanted retrieval but not directly to the ensuing forgetting. Given the cited ev-idence linking the top-down modulation to forgetting (Apšvalka et al. 2022; Benoit & Anderson 2012; Depue et al. 2007; Gagnepain et al. 2014; Hu et al. 2017; Mary et al. 2020; Paz-Alonso et al. 2013), we suggest that this is a rather unlikely account. It certainly would also be less parsimonious than postulating a common process that yields both a transient and a sustained effect.

Second, overall, recall performance was close to ceiling in all conditions, indicating that memories were generally strong in our experiment. This was probably a consequence of the elaborate test-feedback procedure that we had implemented to ensure a sufficient learning rate of the large stimulus set. We might have observed stronger SIF in the M1 condition if the initial encoding strength of the M1 memories had been weaker. In this case, the procedure would have been more powerful for detecting a detrimental effect of the dlPFC stimulation on intentional forgetting.

Third, due to technical constraints and safety regulations, we only applied TMS during 400 ms of each suppression trial. We had carefully chosen our time window to increase the chance that we would disrupt the early stages of the suppression phase. This may have been sufficient to hinder people’s ability to control unwanted memories compared with the M1 condition. However, the remaining time of the trial and the subsequent fixation period may have provided sufficient opportunity to suppress the memories to some degree.

Indeed, our instructions stressed that participants were required not only to prevent the retrieval of the memory but also to push it out of awareness if it came to mind by accident. Accordingly, participants may still have reactively engaged in suppression to purge the intruded memory, even when initial attempts at stopping retrieval failed due to dlPFC stimulation (Benoit et al. 2015; Levy & Anderson 2012). Indeed, although our questionnaire data demonstrated that participants were hindered in their ability to suppress memories, they did not indicate a total inability to keep the memories out of awareness.

Fourth, though the E-field modeling corroborates that our manipulation likely affected dlPFC functioning, we do not suggest that it inflicted a complete functional disruption of this region (Bergmann & Hartwigsen 2021; Hartwigsen & Silvanto 2023; Numssen et al. 2024). The residual functioning of this region may have induced some degree of forgetting– though it was not sufficiently strong to become significant, it may have been strong enough to minimize the difference to the M1 condition.

Taken together, these points suggest that the transient ability to prevent a memory from coming to mind may be more sensitive to the impact of the dlPFC stimulation than suppression-induced forgetting. However, the obtained evidence for a disruption of memory suppression is consistent with other evidence from non-invasive brain stimulation on the contribution of the right dlPFC to active forgetting. For example, transcranial direct current stimulation of the right dlPFC disrupts the phenomenon that targeted retrieval of some memories leads to the incidental forgetting of other,competing memories (Penolazzi et al. 2014; Stramaccia et al. 2017), i.e., retrieval-induced forgetting (Anderson et al. 1994). It also seems to prevent intentional forgetting at encoding (i.e., directed forgetting, Imbernón et al. (2022), Silas & Brandt (2016), Stauch et al. (2020), and Xie et al. (2020)). The current study complements this finding by showing that disrupting the right dlPFC also makes it harder to intentionally control unwanted memories at the retrieval stage.

Intriguingly, the right dlPFC may support an even broader function in initiating inhibitory control. fMRI and EEG studies have provided evidence that a common part of this region is involved in both suppressing memories and suppressing motor actions (Apšvalka et al. 2022; Castiglione et al. 2019; Depue et al. 2010; Depue et al. 2016; Guo et al. 2018).Consistent with this account, evidence from noninvasive brain stimulation has demonstrated that interfering with activity in this region impairs the inhibition of prepotent motor responses (e.g., Hannah et al. (2020), Obeso et al. (2013), and Stramaccia et al. (2015)). This has led to the hypothesis that the right dlPFC may be part of a supramodal inhibitory control mechanism deputed to stop a variety of prepotent responses across different cognitive domains (Wessel & Anderson 2024).

This study provides the first causal evidence for a role of the dlPFC in suppressing unwanted memories. However, future research could be benefit from addressing several key points. First, in light of the current data, it should ideally employ larger sample sizes to achieve sufficient power to detect also more subtle changes in SIF. Second, future studies may more precisely equate the strength of the induced E-field at the respective stimulation sites (Numssen et al. 2024), to further bolster inferences about the spatial specificity of the obtained effects. Third, future research could incorporate trial-by-trial intrusion ratings, to obtain an online measure of memory control. In the present study, we measured intrusion control with a post-experimental questionnaire to avoid additional meta-cognitive task demands. Trial-by-trial ratings of intrusions, however, may be a more sensitive and reliable measure of the disruptions induced by TMS. It would thus be particularly adequate to examine how the transient disruption of dlPFC activity changes the typical association between declining intrusions and long-term forgetting (Legrand et al. 2022; Levy & Anderson 2012).

That being said, it is important to note that including such online ratings may alter the metacognitive structure of the task. In having to reflect on the immediately preceding trial, participants may inadvertently think and thus elaborate on the memory that had just intruded. It may also add to an expectation of a final memory test, which may entice participants to rehearse No-Think memories. If so, these changes in the task might potentially weaken SIF effects. Moreover, responding on each trial via button press may potentially also interact with stimulation.

Finally, future work should build on this study to examine the contributions and interactions of other regions. Of particular interest may be those parietal areas involved in bottom-up attentional capture and top-down processes under-lying the retrieval of memories (Cabeza 2008; Cabeza et al. 2011; Hutchinson et al. 2014; Sestieri et al. 2017).

To conclude, this study provides critical evidence that disruption of the right dlPFC makes it harder to control unwanted memories. It causes a transient disruption of a suppression mechanism, corroborating the hypothesis that this region initiates an inhibitory process that interferes with hippocampal retrieval. However, given the inconclusive results with respect to suppression-induced forgetting, it remains to be determined whether these disruptions can also weaken the sustained after-effects of this process. Nevertheless, the presented data provide the first evidence for the causal contribution of the dlPFC to retrieval suppression.

## ACKNOWLEDGEMENTS

This work as well as DFS, FB, KL, and RGB were funded by a Max Planck Research Group awarded to RGB. ON was supported by the Federal Ministry of Education Germany (Bundesministerium für Bildung und Forschung, BMBF, Grant no. 01GQ2201). GH was supported by the Lise Meitner Excellence Program of the Max Planck Society, the European Research Council (ERC consolidator grant FLEXBRAIN, ERC-COG-2021-101043747) and the German research Foundation (DFG, HA 6314/4-2, HA 6314/10-1).

## Data Availability

All described data and the JASP file are available on the Open Science Framework at https://osf.io/f8m63/?view_only=eb20360279ec47188464caeb2a37cbee.

